# A retrograde GCN2/eIF2α, but ATF4 independent, mechanism maintains synaptic strength under acute amino acid scarcity at the NMJ

**DOI:** 10.1101/2022.11.03.515133

**Authors:** Grant Kauwe, Megumi Mori, Edward H. Liao, Gary Scott, A. Pejmun Haghighi

## Abstract

Neuronal response to nutrient availability plays an important role in the maintenance of cellular homeostasis and behavioral response to the environment in higher eukaryotes. However, we know little about how neuronal function is influenced by acute changes in nutrients at high resolution. Taking advantage of powerful fly genetics and the amenability of the *Drosophila* larval neuromuscular junction (NMJ), we have investigated the synaptic response to acute amino acid restriction. Our findings indicate that the presence of general control nonderepressible 2 (GCN2) and phosphorylation of its target eukaryotic initiation factor 2 alpha (eIF2α) are essential for the ability of the NMJ to maintain normal neurotransmitter output when the larvae are deprived of amino acids. Surprisingly, activating transcription factor 4 (ATF4), which normally acts downstream of GCN2/eIF2α, appears dispensable in this regulation. Furthermore, we show that GCN2/eIF2α dependent cascade acts retrogradely from muscle back to motoneuron to adjust synaptic release. These results provide a mechanistic insight into the intricate regulation of synaptic strength through the action of GCN2 when organisms are faced with amino acid scarcity.

## INTRODUCTION

Many nutrient sensing mechanisms work in concert to allow for appropriate cellular processes as organisms are faced with conditions of food scarcity or abundance^1^. In higher eukaryotes with more complex nervous systems, it has been appreciated that neuronal control plays an important role in sensing changes in nutrient availability as well as in orchestrating an organismal wide response to these changes^2,3^. Regulation of cap-dependent translation by the target of rapamycin (TOR) and limiting the availability of the ternary complex through phosphorylation of eukaryotic initiation factor 2 alpha (eIF2α) by general control nonderepressible 2 (GCN2) are among the best studied cellular mechanisms that are triggered in the response to amino acid scarcity from yeast to mammals; interestingly, both of these pathways have also been implicated in the regulation of synaptic plasticity^1–7^. However, existing knowledge lacks many details about how these pathways control basal synaptic activity in response to acute changes in nutrients.

Phosphorylation of eIF2α by GCN2 is part of a conserved cellular response to stress in organisms, which is also known as the integrated stress response (ISR). Once phosphorylated, eIF2α curtails the formation of the 43S preinitiation complex, leading to a slowing of the rate of translation initiation^4,8^. In addition to GCN2, eIF2α is also phosphorylated by PERK in response to ER stress and accumulation of unfolded proteins, by PKR in response to viral infection, and by HRI in response to Heme deficiency in mammals ^9,10^. Phosphorylation of eIF2α limits the availability of the ternary complex, critical for translation initiation, and thus dampens the overall rate of translation in cells^8^. Paradoxically, a small number of mRNAs with highly structured 5’UTRs, of which activating transcription factor 4 (ATF4) is the archetype, are better translated in response to eIF2α phosphorylation^11,12^. ATF4 plays a critical role in mediating the downstream effect of eIF2α phosphorylation by transcriptionally activating a number of stress response genes^13^.

We have taken advantage of the powerful genetic approaches in Drosophila and the amenability of the Drosophila larval neuromuscular junction (NMJ) to investigate the effect of acute amino acid (AA) deprivation on evoked synaptic release at a single NMJ. Our findings indicate that GCN2 is critical in postsynaptic muscles, but not in motoneurons, to ensure that synaptic strength is maintained under AA restriction. We further show that, as expected, eIF2α phosphorylation by GCN2 is essential; however, our genetic analysis indicates that ATF4 is surprisingly dispensable. Therefore, our results point to a new mode of stress response at the synapse that is critical for the maintenance of synaptic strength under acute amino acid restriction.

## RESULTS/DISCUSSION

### Synaptic strength critically depends on GCN2 under amino acid deprivation

Glutamatergic synapses at the *Drosophila* larval neuromuscular junction (NMJ) employ a wide number of mechanisms to tightly regulate synaptic strength. The accessible anatomy of the larval neuromuscular innervation allows for the interrogation of synaptic activity at a single identifiable NMJ using intracellular electrophysiological methods^7^. We asked whether restricting amino acids in food would influence the basal synaptic transmission at the NMJ. We found that these synapses show remarkable capacity for maintaining basal synaptic strength under conditions of amino acid scarcity. When larvae are acutely (6 hrs.) fed on AA-restricted food, the amplitude of the miniature excitatory postsynaptic currents (mEPSCs), evoked postsynaptic currents (EPSCs) and quantal content (QC) remain unaffected (Figure 1a and 1c). One of the main cellular responses to amino acid scarcity is the activation of the integrated stress response kinase General Control Nonderepressible 2 (GCN2).

**Figure 1.**
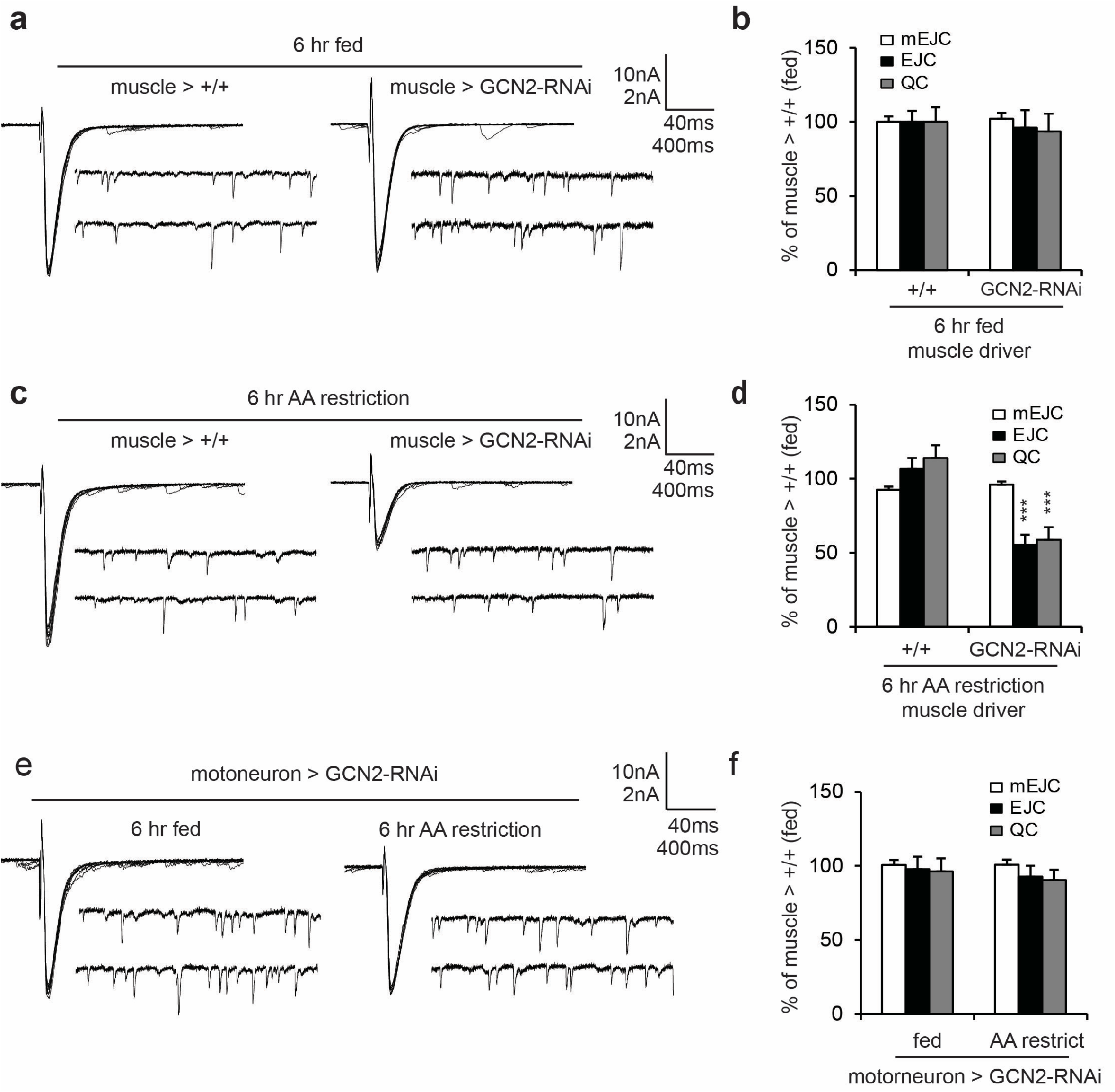
GCN2 is required for NMJ homeostasis in response to AA-restriction. **a)**, **c)** and **e)** are representative traces of miniature excitatory junctional currents (mEJC) and evoked excitatory junctional currents (EJC) recorded from muscle 6, third instar larvae under normal (fed) or amino acid restriction (AA restriction) conditions as indicated. MHC-Gal4 and BG380-Gal4 are the muscle and neuronal drivers, respectively. **b)**, **d)** and **f)** are quantification of average mEJC, EJC and quantal content corresponding to genotypes in a, c and e, respectively. Statistics are paired wise t-Tests. ** is *p*<0.01 and *** is *p*<0.001.

We, therefore, asked whether activation of GCN2 is required for maintaining basal synaptic strength at the NMJ. When larvae were raised on normal food, knockdown of GCN2 in either postsynaptic muscles or presynaptic motoneurons had no effect on baseline electrophysiological measurements (Figure 1a, b, e and f). However, on AA-restricted food, transgenic knock-down of GCN2 (Supplementary Figure 1a) in postsynaptic muscles, but not presynaptic motoneurons, led to a severe reduction in synaptic function, such that the baseline EPSC and QC dropped by more than 40% (Figure 1c-f). We found that the requirement for postsynaptic GCN2 remained under more prolonged (24hr) amino acid scarcity, suggesting that no adaptation is occurred in the absence of GCN2 at least for 24 hrs. (Supplementary Figure 1d, e). We did not detect any changes in the average amplitude of mEPSCs, suggesting that the changes in quantal content are due to presynaptic changes in neurotransmitter release. Next, to assess whether GCN2 knockdown could have had a significant effect on synaptic structures and NMJ growth, we examined NMJ morphology. GCN2 knockdown did not affect the number of synaptic boutons (Supplementary Figure 1b, c).

Together, these results demonstrate a critical role for GCN2 in the maintenance of synaptic strength at the NMJ under conditions of amino acid scarcity and indicate that GCN2 exerts its function in a retrograde manner from muscle to motoneuron.

### eIF2α phosphorylation is critical for the maintenance of synaptic strength under amino acid deprivation

Under conditions of amino acid scarcity, GCN2 phosphorylates its target eIF2α and thereby curtails the formation of the 43S preinitiation complex which leads to a slowing of the rate of translation initiation^4,8^. Consistent with activation of GCN2 under amino acid deprived conditions, we detected a strong increase in eIF2α phosphorylation in larval muscle preparations (Figure 2a-c). We found that larvae feeding on AA-restricted food showed a significant increase in eIF2α phosphorylation compared to larvae feeding on standard food, as evidenced by western blot analysis of larval muscle extracts (Figure 2b, c). If increased phosphorylation of eIF2α is required for the maintenance of the baseline synaptic function under AA restriction, then preventing eIF2α phosphorylation should mimic the effect of loss of GCN2. GCN2 phosphorylates eIF2α at the serine residue at position 51 in mammals and yeast, and position 50 in *Drosophila.*

**Figure 2.**
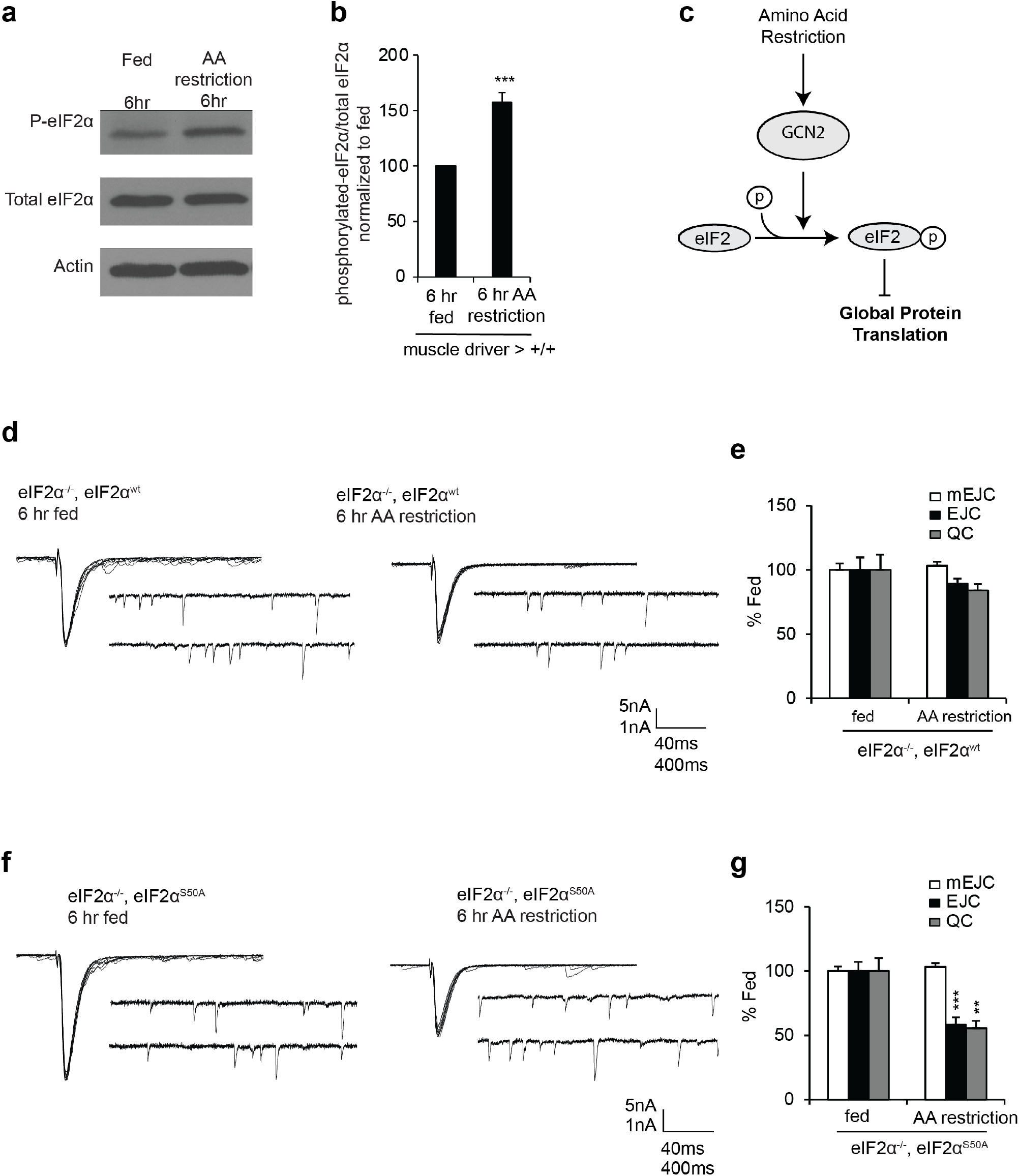
eIF2α is required for NMJ homeostasis in response to AA-restriction. **a)** Amino acid restriction leads to enhancement in eIF2α phosphorylation. **b)** Western blot using dissected larval muscle fillet preparations. **c)** Quantification of three western blots. **d)** Larvae expressing a genomic transgenic insert carrying the genomic region of eIF2α (eIF2α^WT^) or **f)** expressing the same genomic locus plus a mutation for the eIF2α phosphorylation site by GCN2 (S50A in Drosophila) (eIF2α^S50^) in eIF2α mutant background. Base line electrophysiological properties in larvae carrying the S50A mutation are indistinguishable from WT counterparts, but they fail to maintain quantal release under amino acid restricted conditions. **e)** and **g)** are quantification of electrophysiological recordings in d and f, respectively.

To test the relevance of eIF2α phosphorylation, we generated transgenes containing the complete eIF2α gene with S50 intact or substituted by an alanine, eIF2α^WT^ and eIF2α^S50A^, respectively. We combined either transgene with an eIF2α KO mutant, eIF2α^G0272^, that is otherwise adult homozygous lethal^7^. We were able to rescue the reduction in the amplitude of mEJCs in hemizygous male or homozygous female eIF2α^G0272^ mutant larvae such that the average mEJC amplitudes were indistinguishable from control larvae under normal food conditions (Figure 1a and 2d). Interestingly however, when these larvae were placed on AA-restricted food, base line synaptic strength was no longer maintained in larvae expressing eIF2α^S50A^ transgene, demonstrating that phosphorylation of eIF2α at S50 is essential for the ability of larval NMJ to maintain its synaptic strength under AA-restricted conditions (Figure 2f, g).

### ATF4 Drosophila homologue, CRC, is not required for maintenance of synaptic function under AA-restriction

Since ATF4 acts as a master regulator that activates a number of genes that orchestrate the cellular stress response^13^, we hypothesized that ATF4 is required downstream of GCN2 activation and phosphorylation of eIF2α, initiate the retrograde signaling mechanism required for the maintenance of NMJ synaptic strength under AA-restricted conditions. To test this idea, we genetically knocked down the *Drosophila* homologue of ATF4, Crc, in postsynaptic muscles (Figure 3a). Surprisingly, knockdown of Crc under AA-restricted conditions did not phenocopy either GCN2 knockdown or inhibition of eIF2α phosphorylation (Figure 3b). Both spontaneous or evoked synaptic responses were not changed in response to postsynaptic knockdown of Crc under normally fed or AA-restricted conditions up to 24hrs (Figure 3c). These results suggest that maintenance of synaptic activity under AA-restricted conditions does not require upregulation of ATF4 and the canonical activation of the integrated stress response at the Drosophila larval NMJ.

**Figure 3.**
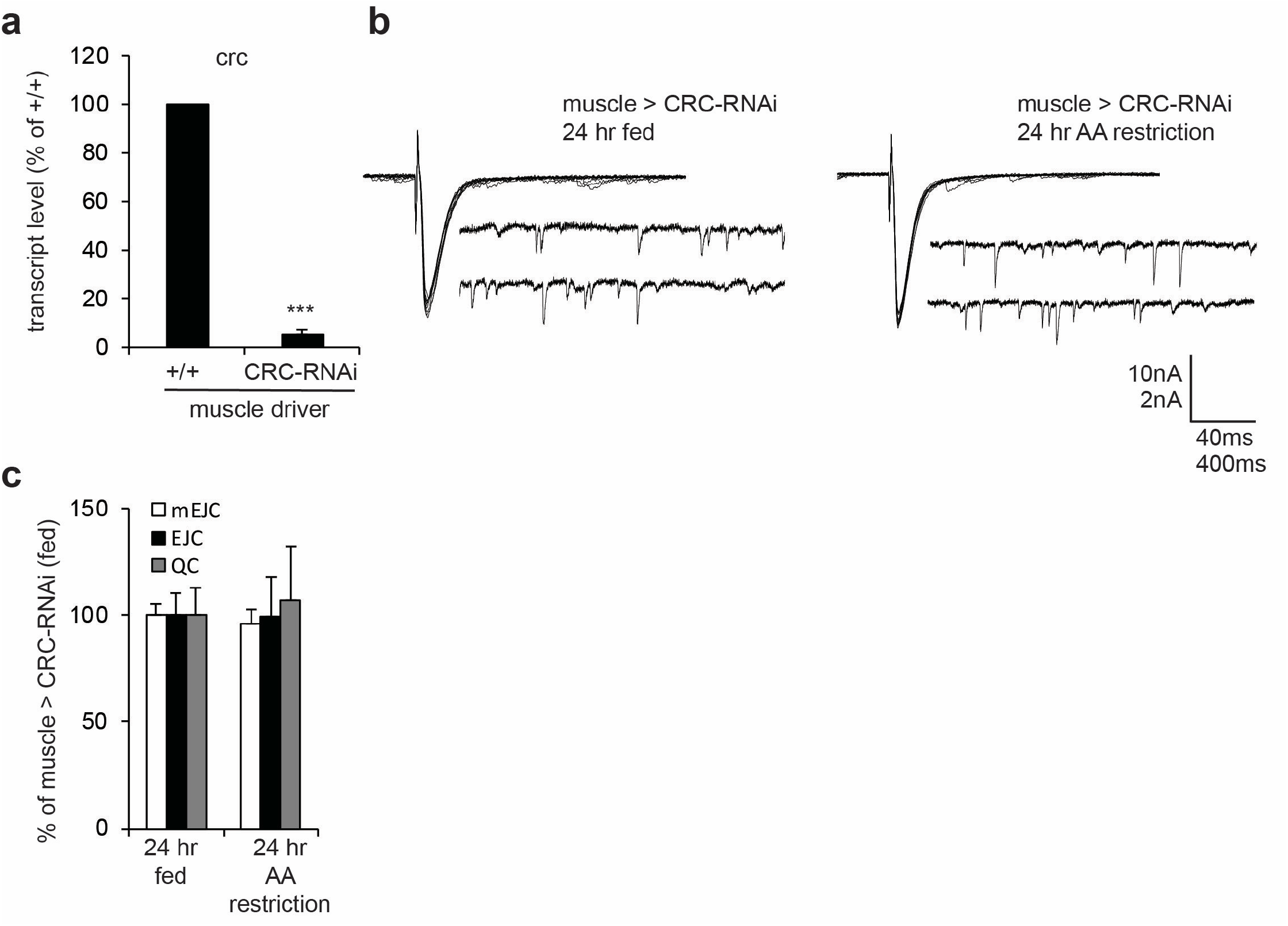
ATF4 *Drosophila* homologue, CRC, is not required for maintenance of synaptic function under AA-restriction. **a)** Transgenic knockdown of Crc, the fly homologue of ATF4, in postsynaptic muscles effectively reduces its transcript level. **b)** Postsynaptic knockdown of Crc does not influence synaptic strength at the larval NMJ under amino acid deficient conditions. **c)** Quantification of the results in b. No statistical significance was observed.

Our results together build a strong case for the existence of a retrograde GCN2/eIF2α dependent mechanism that is triggered in postsynaptic muscles under AA-restricted conditions. Our findings suggest that as a result of eIF2α phosphorylation, a cellular program required for the maintenance of synaptic activity under amino acid scarcity is triggered in an ATF4 independent manner. Based on the conserved nature of this signaling pathway^13–18^, it is likely that a similar mechanism is at work at the vertebrate NMJ or potentially at higher CNS synapses. It will be of great interest to identify translational target(s) of GCN2/p-eIF2α in postsynaptic muscles. We predict that these target genes are likely to have a complex 5’-UTR that contain upstream open reading frames similar to those present in the 5’-UTR of ATF4/Crc^19–22^.

## Materials and Methods

### Fly Genetics

Flies and larval progenies were maintained at 25°C incubators on a 12-hour light/dark cycle on standard medium. Third instar larvae were collected for analysis. See Fly Stock List for a comprehensive list of fly stocks used.

|  |  |  |
| --- | --- | --- |
| *D. melanogaster*. RNAi against GCN2. y[1] sc[*] v[1] sev[21]; P{y[+t7.7] v[+t1.8]=TRiP.GL00267}attP2 | Bloomington Drosophila Stock Center | BDSC: 35355 |
| *D. melanogaster*. RNAi against GCN2. y[1] sc[*] v[1] sev[21]; P{y[+t7.7] v[+t1.8]=TRiP.GL00267}attP2 | Bloomington Drosophila Stock Center | BDSC: 35355 |
| *D. melanogaster*. Muscle driver MHC-Gal4 | (Schuster et al., 1996) |  |
| *D. melanogaster*. Muscle driver. 24B-Gal4 | (Brand & Perrimon, 1993) |  |
| *D. melanogaster*. RNAi for dnc. y[1] v[1]; P{y[+t7.7] v[+t1.8]=TRiP.JF02561}attP2 | Bloomington Drosophila Stock Center | BDSC: 27250 |
| *D. melanogaster*. RNAi for mfr. y[1] sc[*] v[1] sev[21]; P{y[+t7.7] v[+t1.8]=TRiP.GLV21054}attP2 | Bloomington Drosophila Stock Center | BDSC: 35689 |
| *D. melanogaster*. RNAi for mtt. y[1] sc[*] v[1] sev[21]; P{y[+t7.7] v[+t1.8]=TRiP.HMS02793}attP40 | Bloomington Drosophila Stock Center | BDSC: 44076 |
| *D. melanogaster*. RNAi for Rab26. y[1] v[1]; P{y[+t7.7] v[+t1.8]=TRiP.JF01684}attP2 | Bloomington Drosophila Stock Center | BDSC: 31177 |
| *D. melanogaster*. RNAi for Rph. y[1] v[1]; P{y[+t7.7] v[+t1.8]=TRiP.JF01970}attP2 | Bloomington Drosophila Stock Center | BDSC: 25950 |

### Acute amino acid restriction experiments

Larvae were transferred to control food (1L water, 13.8g bacteriological agar, 50g sucrose, 80g corn flour, 24g brewer’s yeast, 0.5g methylparaben, 5mL EtOH, and 3mL propionic acid) or amino acid restricted food, which is exactly the same without addition of Yeast.

### Western Blot

Wandering third instar larvae were dissected in cold PBS and the muscles were collected. 10 larvae were used per biological replicate for a total of 3 biological replicates. Western blot was performed according to standard protocols. The following primary antibodies and concentrations were used: anti-phospho-eIF2α 1:1000 (S371, 119A11, Cell Signaling), anti-non-phospho-eIF2α 1:1000 (EIF2S1, ab26197, Abcam), anti-Actin 1:5000 (C4, MAB1501, Sigma-Aldrich). The following secondary antibodies and concentrations were used: HRP-conjugated anti-mouse IgG, IgM 1:2000 (A-10677, Invitrogen), HRP-conjugated anti-rabbit IgG, IgM 1:2000 (G21234, Invitrogen).

### Quantitative PCR

Total RNA was extracted from 5 wandering third instar larvae with standard Trizol extraction. cDNA was produced using the BioRad iScript cDNA synthesis kit (Cat No. 170-8891). qPCR data was generated using the BioRad iTaq Universal SYBR Green Supermix on the BioRad CFX96 detection system.

### Confocal imaging and image analysis

Muscle 6/7 from segment A3 of third instar larvae were dissected as previously described. Briefly, third instar larvae were dissected in cold HL3 and fixed with 4% Paraformaldehyde in PBS (Phosphate buffered saline) for 10min. Larval fillets were then washed with PBS three times, permeabilized and blocked with 5% Normal Goat Serum (NGS) in PBT (PBS with 0.1% Triton X-100) for 1hr and placed in primary antibody overnight at 4°C. The larvae were then washed three times for 15min in PBT, placed in secondary antibody for 2hrs, washed three times for 15min with PBT and mounted in Vectashield (Vector labs). The following antibodies and concentrations were used for quantifying the boutons at the NMJ: anti-Dlg 1:500 (4F3E3E9, Developmental Studies Hybridoma Bank), Cy3-conjugated goat anti-HRP 1:250 (123-165-021, Jackson ImmunoResearch). The number of boutons were quantified using confocal images acquired on LSM700 with 40x objectives.

### Electrophysiology

Standard two-electrode electrode voltage clamp technique was used in dissected third instar larvae in HL3 with 0.5 mM Ca++ (Penney et al., 2012).

### Statistical Analysis

All statistical analyses were performed using GraphPad Prism software or RStudio (RStudio). Further statistical details for each experiment can be found in the corresponding figure legend. Mean ± s.e.m. is shown for quantifications. Asterisk P-values are as follows: *<0.05, **<0.01, ***<0.001.

## ACKNOWLEDGEMENTS

We thank N. Sonenberg, S. Sigrist and for providing us with reagents. We thank all the present and past members of the Haghighi lab for helpful discussions about the manuscript, particularly Ammar Aly for help with edits and M. Moujahidine for administrative support.

Images were collected at the Buck Institute Imaging and Morphology Core. We thank the Bloomington Center and Vienna *Drosophila* Resource Center for fly stocks and the University of Iowa Developmental Studies Hybridoma Bank for antibodies. This work was supported by a T32 fellowship to G.K., and an NIH grant (NS R01 NS082793) to APH.

## AUTHOR CONTRIBUTIONS

G.K., M.M., E.H.L. and A.P.H. designed the experiments. G.K., M.M., E.H.L. and G.A., performed the experiments. M.M. and A.P.H. prepared the manuscript.

## MATERIALS AND CORRESPONDENCE

Correspondence and requests for materials should be addressed to A. Pejmun Haghighi.

