## Supplemental Figures for "A retrograde GCN2/eIF2α, but ATF4 independent, mechanism maintains synaptic strength under acute amino acid scarcity at the NMJ"

**Supplementary Figure 1. GCN2 is dispensable for synaptic structural growth**

**a)** transgenic RNAi effectively reduced GCN2 expression when expressed in postsynaptic muscles using MHC-Gal4 driver.

**b)** Confocal images of NMJs at muscles 6/7 in third instar larvae (third abdominal segment) stained with postsynaptic density marker (discs large, Dlg in green) and presynaptic marker Hrp in red.

**c)** Number of boutons are not affected as a result of GCN2 knockdown.

**d)** Electrophysiological recordings from muscle 6 in third abdominal segments from control larvae or larvae overexpressing GCN2-RNAi in muscle under aa restricted condition for 24hrs.

**e)** Quantification of Miniature synaptic currents, evoked synaptic currents and quantal content corresponding to data in d. ** is P<0.01, *** is P<0.001, Student-t tests were performed.

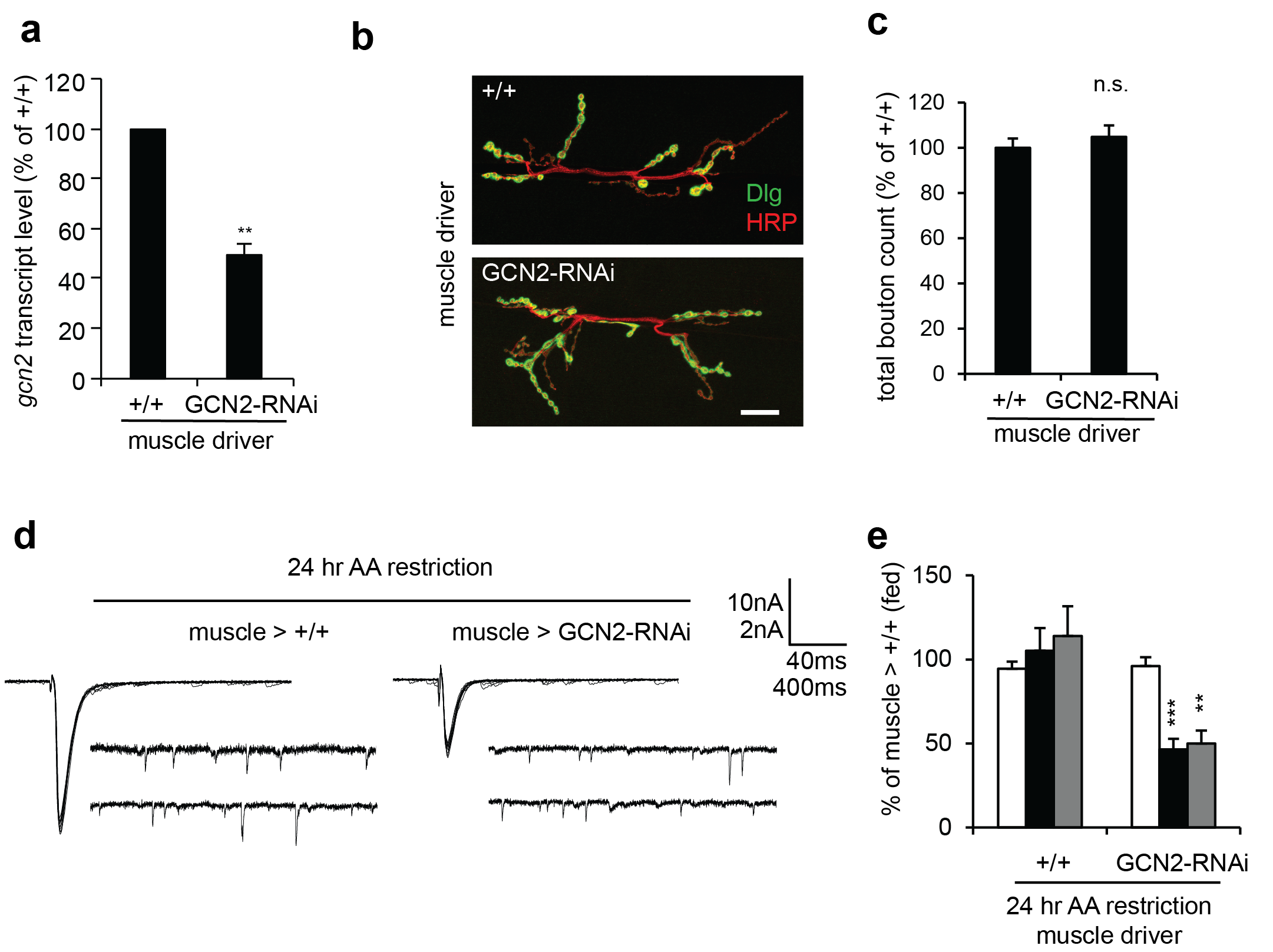

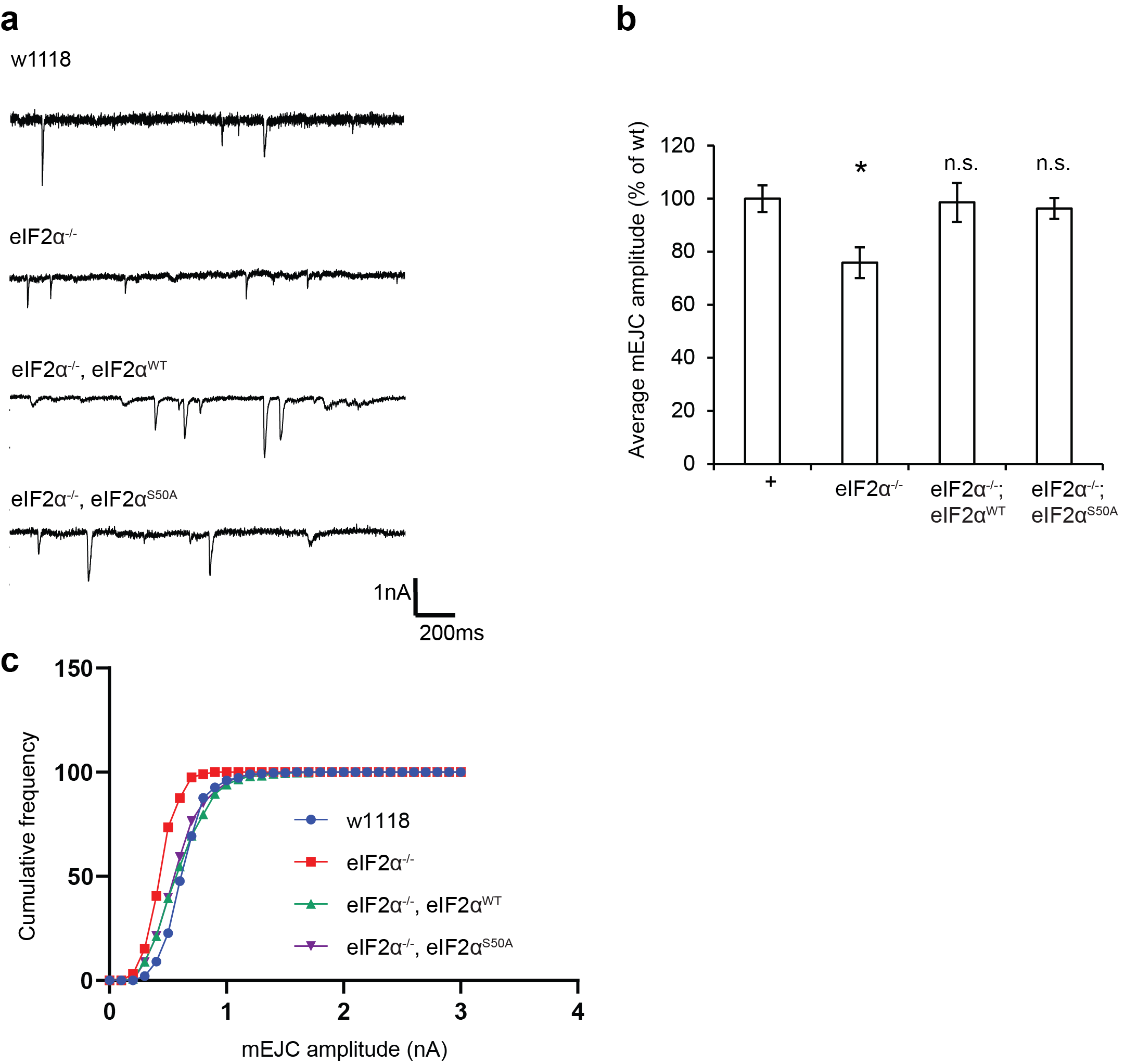

**Supplementary Figure 2. WT and S50A transgenes restore mEJC amplitude in eIF2α mutant larvae.**

**a)** Sample mEJCs from the respective genotypes.

**b)** Quantification of mEJC average amplitude corresponding to data in a). * is P<0.05, Student-t test was performed.

**c)** Cumulative distribution of miniature synaptic amplitudes. Both transgenes effectively restore normal distribution.
